# A ClyA nanopore tweezer for analysis of functional states of protein-ligand interactions

**DOI:** 10.1101/727503

**Authors:** Xin Li, Kuohao Lee, Jianhan Chen, Min Chen

**Affiliations:** Department of Chemistry, University of Massachusetts Amherst, Amherst, Massachusetts 01003, USA; Molecular and Cellular Biology Program, University of Massachusetts Amherst, Amherst, Massachusetts 01003, USA; Department of Biochemistry and Molecular Biology, University of Massachusetts Amherst, Amherst, Massachusetts 01003, USA

## Abstract

Conformational changes of proteins are essential to their functions. Yet it remains challenging to measure the amplitudes and timescales of protein motions. Here we show that the ClyA nanopore can be used as a molecular tweezer to trap a single maltose-binding protein (MBP) within its lumen, which allows conformation changes to be monitored as electrical current fluctuations in real time. The current measurements revealed three distinct ligand-bound states for MBP in the presence of reducing saccharides. Our biochemical and kinetic analysis reveal that these three states represented MBP bound to different isomers of reducing sugars. These findings shed light on the mechanism of substrate recognition by MBP and illustrate that the nanopore tweezer is a powerful, label-free, single-molecule approach for studying protein conformational dynamics under functional conditions.

---

Proteins are dynamic entities that sample multiple conformations over a range of timescales^1,2^. Understanding how proteins work requires a full description of their conformational dynamics over time. Time-dependent and transient protein dynamics are challenging to observe with structural approaches such as X-ray crystallography and electron microscope. Nuclear magnetic resonance (NMR) spectroscopy can measure protein dynamics at multiple time scales^2,3^, but its application is laborious and limited to relatively small proteins that can be stably prepared at high concentrations. As an ensemble method, NMR also lacks the ability to resolve multiple weakly populated conformational states and conformational heterogeneities^4^. In contrast, single-molecule techniques such as fluorescence resonance energy transfer (smFRET) are powerful tools to interrogate protein dynamics^5–7^. The single molecule resolution allows the revealing of the previously masked conformational dynamics^8^ and heterogeneities in enzymatic function^9,10^, as well as transient intermediates and stochastic events^11^. However, due to photobleaching, blinking and limited photon flux of the fluorophores, smFRET has difficulty to resolve the very short-lived states or monitor the conformational variability of the same molecule over a long duration^12^. Recently, methods based on electrical measurements, which do not have the limitations associated with fluorophores, have been explored to expand single-molecule analyses for recording transient motions and intermediate states of proteins^13^.

Nanopores have emerged as a powerful, single-molecule analytical tool to address fundamental problems in chemistry and biology^6,12,14^. A nanopore detection platform is composed of a single hole in an insulating membrane that separates two electrolyte chambers. An analyte molecule interacting with the nanopore occludes the pore and causes a transient change in ionic current. Taking advantage of the high temporal resolution (μs-ms) of the nanopore measurement, a biological nanopore Cytolysin A (ClyA) from *Salmonella typhi* has been applied to monitor fast action of substrate binding, or enzyme activities by confining a single protein within its lumen^15,16^. Current measurements from a ClyA nanopore from *Escherichia coli* have been shown to distinguish apo avidin from biotin-bound avidin, even though the root mean square deviation (RMSD) between the C_α_ atoms of the two states is only 0.42 Å^17,18^; this demonstrates that the nanopore approach has high spatial sensitivity and the potential to detect subtle differences between protein conformers.

MBP is a periplasmic binding protein responsible for the uptake and efficient catabolism of maltodextrins in *E. coli*. MBP is ellipsoidal and composed of two distinct globular domains with a ligand binding site located at the hinge region of two domains^19–22^ (Fig. 1a). Apo MBP adopts an open conformation where the groove between the domains is readily accessible to solvents and ligands^23^. In the ligand bound state, MBP becomes more compact with the two domains moving towards each other by ∼1 nm^23^ (Fig. 1a) The large-scale domain motion triggered by ligand-binding makes MBP a suitable model for studying the conformation dynamics by both ensemble and single molecule approaches, such as NMR and smFRET^12,24,25^.

**Figure 1.**
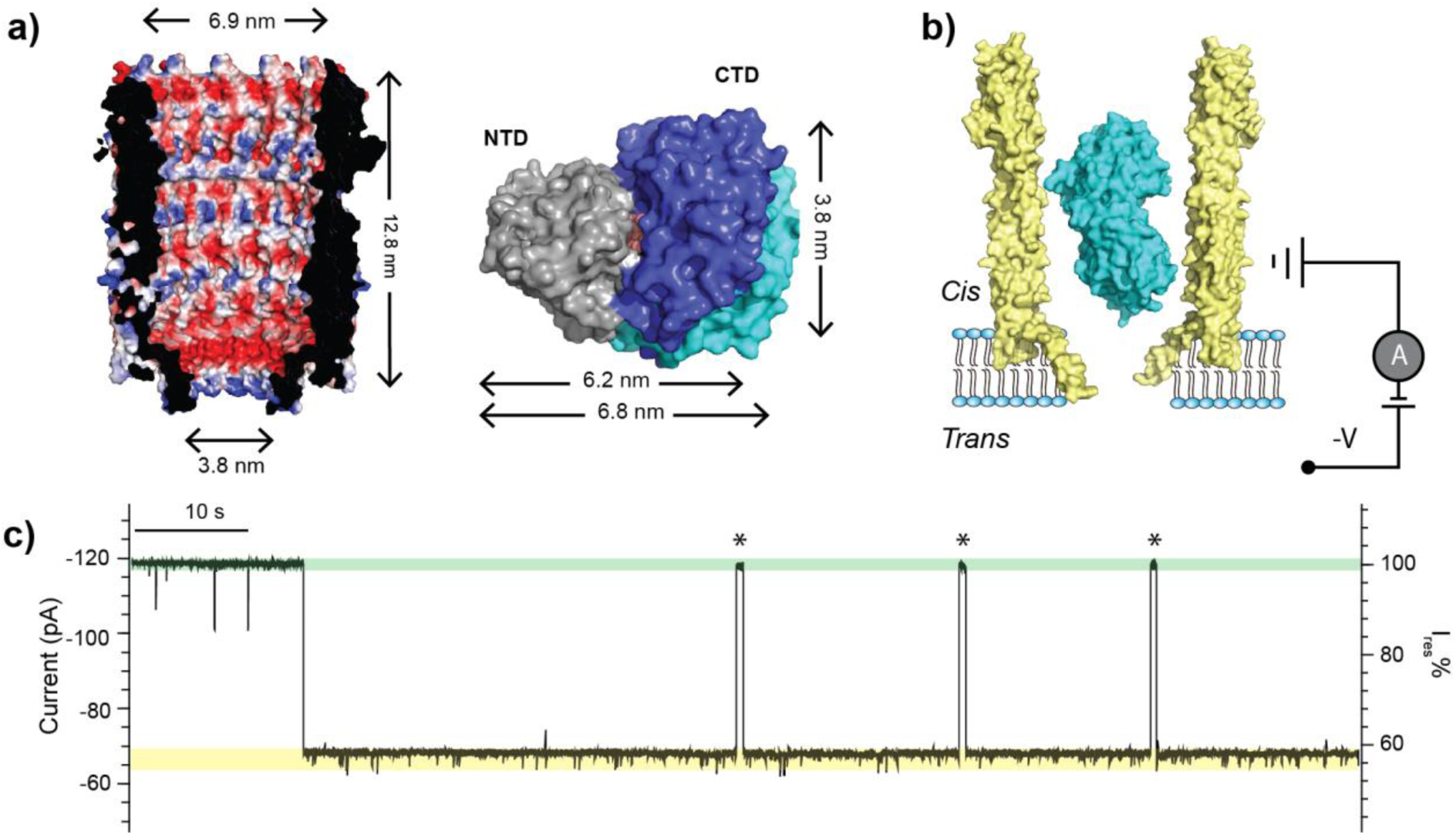
Trapping of MBP within the ClyA nanopore. (a) A cross-section view of the structure of the ClyA nanopore (PDB:2WCD) showing the electrostatic potential of its inner surface (left). Surface representation of MBP structures in the ligand-free (PDB:1JW4) and bound (PDB:1ANF) states (right). The two structures are aligned by the N-terminal domain (NTD, grey), with the C-terminal domains (CTD) shown in cyan and blue for the apo and holo states, respectively. The bound maltose is shown in orange. (b) Schematic representation of an MBP trapped in ClyA with negative potential applied on the *trans* side. (c) Representative current trace of ClyA in the presence of MBP. The open pore current is highlighted in green, while the blockades caused by MBP in yellow. Asterisks represent MBP escape events from the nanopore. The current traces were collected at −80 mV in the buffer 150 mM NaCl, 15 mM Tris-HCl, pH 7.5 in the presence of 56 nM MBP.

In this work, we apply the ClyA nanopore as a molecular tweezer to trap a single maltose-binding protein (MBP) and monitor its conformational changes during ligand binding. Our single-molecule recordings from the ClyA nanopore tweezer revealed the existence of three distinct ligand-bound states that has not been previously recognized. Combining molecule dynamic simulations and current measurements, we determine that these bound states are induced by the α- and β-isomers of maltooligosaccharides and quantify the binding affinities and kinetic rates of all complexes. Our findings shed new lights on the mechanism of substrate recognition by MBP and provide a new label-free framework for deploying the ClyA trapping approach to study the single-molecule protein dynamics in various biological processes.

## Confinement of a single MBP inside the ClyA nanopore

For single channel recording studies, ClyA pores were obtained by incubating ClyA monomers with *n*-dodecyl-β-maltoside^19^. The pores were then added to the *cis* chamber of a recording apparatus. The resulting ClyA pores varied widely in conductance, which suggests that the pores are formed from oligomers of different stoichiometries and sizes^19^. In this work, we chose to work with ClyA pores with a conductance of approximately 1.5 nS, which was the major population and appeared to be the most suitable for trapping MBP. The structure of the *E. coli* ClyA pore can be approximated as a conical frustum with an opening of 6.9 nm in diameter at the top and 3.8 nm at the bottom (Fig. 1a). Therefore, ClyA nanopores can accommodate a single MBP protein (∼3.0 × 4.0 × 6.5 nm^3^; see Fig. 1a) without it being translocated through. ClyA nanopores insert into lipid bilayers in a unidirectional manner due to the large soluble domain, with the nanopore lumen facing the *cis* chamber where both ClyA and MBP were added. When a negative voltage was applied to the *trans* side, current blockades were observed (Fig. 1b), indicating that an MBP was trapped in the pore. When a positive voltage was applied to the *trans* side, ClyA did not capture MBP. The inner surface of ClyA is predominantly negatively charged (Fig. 1a). At pH 7.5, MBP is slightly negatively charged, which suggests that the electroosmotic force is dominant to drive MBP against the electrophoretic force. The duration of MBP trapping is voltage dependent, with higher voltages leading to longer MBP trapping times (Supplementary Fig. 1). Appling voltages higher than −80mV caused self-closure or gating of the ClyA nanopore; therefore, we applied a voltage of −80 mV throughout this study. MBP trapping induced current blockades with a residual current (I_res_ %) of 57.1 ± 1.3 % (n = 45, Fig. 1c). By measuring the dwell time (τ_off_ = 178.0 ± 27.1 s, Supplementary Fig. 1) and inter-event interval (τ_on_ = 1.0 ± 0.1 s, Supplementary Fig. 1), we obtained a capture rate of (17.9 ± 1.8) μM^−1^ s^−1^ and an apparent affinity constant of (3.1 ± 0.6) × 10^−10^ M for the trapping of MBP within the ClyA nanopore.

## Detecting substrate binding to the trapped MBP

ClyA nanopores with trapped MBPs exhibited two types of current signals: one quiet (∼60 % of the time) and one noisier with a smaller residual current (∼40% of the time; Supplementary Fig. 2). We surmised that these two types of signals were induced by MBP that was trapped in ClyA in opposite orientations (Supplementary Fig. 2). Previous research has shown that proteins can adopt different orientations in nanopores^15^. MBP is too large to change orientation while it is inside the nanopore. Calculated by using standard CHARMM charges, both MBP and ClyA appear to have large net dipole moments (∼694.3 D for holo MBP, 773.7 D for apo MBP, and 7580.7 D for ClyA). The dipole-dipole interaction favors the N-terminal domain–up orientation by ∼0.3 kcal/mol under the experimental conditions, which is consistent with the observed percentages of quiet and noisy traces (Supplementary Fig. 2). We focused our analyses on MBPs trapped in the quiet, N-terminal domain–up orientation, because the resulting signals allowed more reliable data analysis.

We observed that the apo MBP populated two states: state 0, a lower blockage state (57.1 ± 1.3 %, n= 45) and state 1, a higher blockage state (54.9 ± 1.5 %, n= 39) (Fig. 2a). State 0 occupied the majority of the recording time: 98.2 ± 2.4 %. These results may indicate spontaneous conformational fluctuations of MBP in the apo state and are consistent with NMR relaxation analyses showing that apo MBP can sample a “hidden” excited state^20^ with a population of ∼5 %.. These results suggest that the nanopore tweezer method can resolve relatively subtle spontaneous protein conformational transitions.

**Figure 2.**
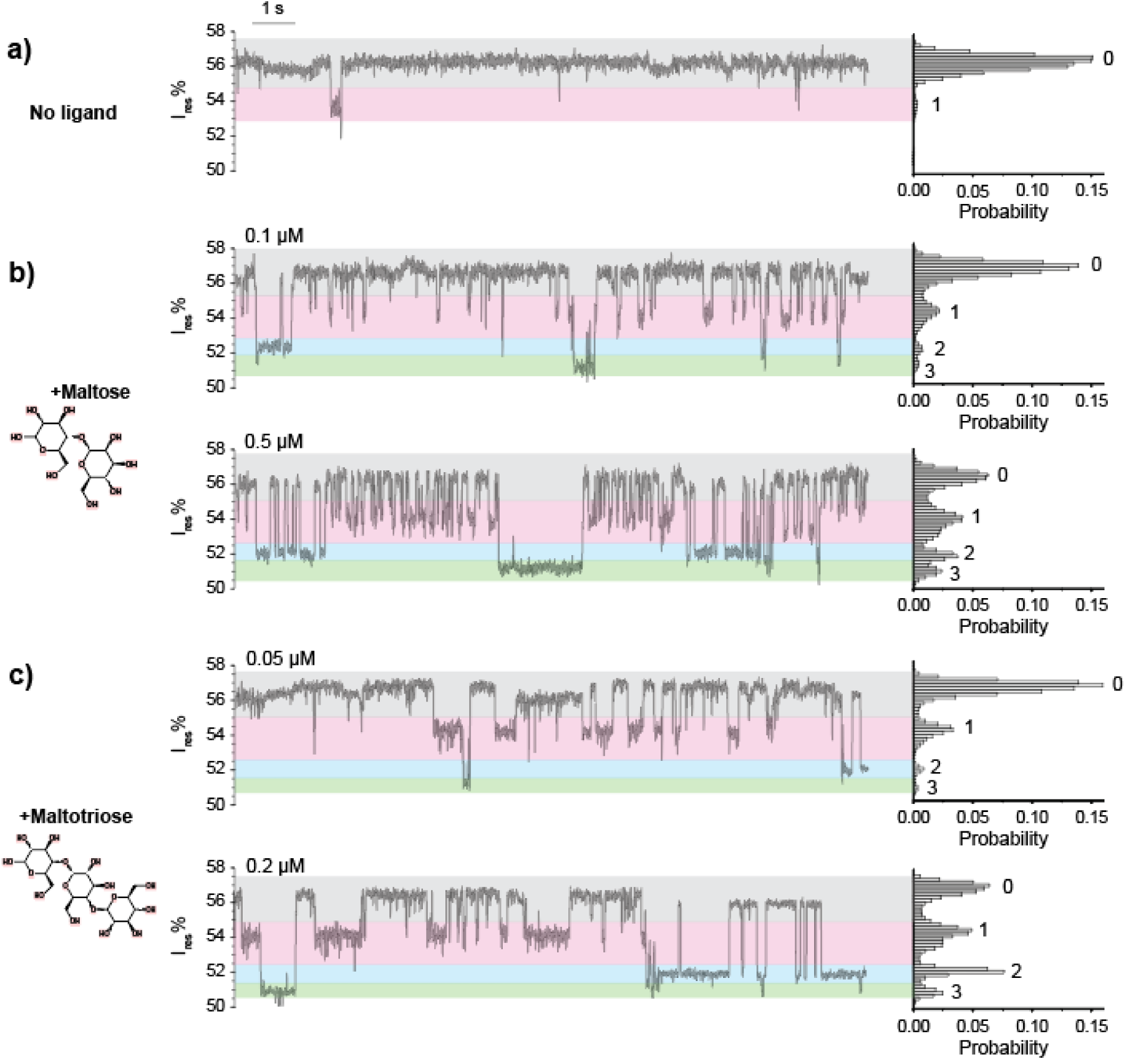
ClyA nanopore current responses induced by ligand interactions with trapped MBP. (a-c) Typical traces and corresponding histograms of ClyA pores containing a trapped MBP in the absence (a) and presence of ligand (b, c). The concentrations of maltose (b) and maltotriose (c) are indicated in the figure. All the measurements were performed in 150 mM NaCl, 15 mM Tris-HCl, pH 7.5 at −80mV by applying a Bessel low-pass filter with a 2 kHz cutoff and sampled at 5 kHz.

## Detection of MBP-ligand interaction within a nanopore

MBP can bind specifically with maltose, maltodextrins, and cyclodextrins^21,22^. To investigate if the nanopore can detect MBP/ligand binding, we added maltose or maltotriose to the recording chamber in various concentrations. In the presence of maltose, four major current states were observed: state 0, with a residual current of 56.5 ± 0.4 %; state 1, with a residual current of 54.0 ± 0.4 %; state 2, with a residual current of 52.0 ± 0.4 %; and state 3, with a residual current of 51.1 ± 0.4 % (Fig. 2b). Maltotriose induced four states that were almost identical to the states seen with maltose. Raising the maltose or maltotriose concentration increased the frequencies of states 1, 2, and 3 (Fig. 2b&c, Supplementary Fig. 3); this suggests that states 1, 2, and 3 represent the bound states of MBP. To exclude the possibility that the signal was caused by nonspecific binding of sugar molecules to the ClyA nanopore itself, we repeated the experiment without MBP and detected no changes in the current recording (data not shown). To further confirm that the signal was caused by specific ligand binding to the trapped MBP, we added sucrose, a non-substrate disaccharide, into the *cis* chamber and observed no signal change (Supplementary Fig. 4). Together, these results demonstrate that the ligand binding of MBP can be monitored by ClyA nanopore current recording at the single-molecule level in real time.

## Molecular Dynamic Modeling and Simulation

X-ray crystallography shows that ligand-free MBP adopts a more open conformation than the ligand-bound MBP^23^. Therefore, it would be reasonable to expect that apo MBP would cause a greater blockage and thus a lower residual current than bound MBP. However, the nanopore measurements showed that MBP bound to maltose or maltotriose blocked the nanopore less than the apo MBP did. We explored this unexpected result by performing molecular dynamics simulations to characterize how MBP interacts with ClyA.

As expected, the modeling results showed that the membrane-embedded region of ClyA (from −15 to 15 Å in z-depth with 0 defined as the center of membrane) has the smallest solvated ion–accessible cross-section and gives rise to the greatest unit resistance, with or without trapped MBP (Fig. 3c). However, trapped MBP forms additional constrictions above the membrane-embedded region (z-depth > 15 Å) (Fig. 3b&c). These constricted regions have cross-sections comparable to the narrowest portion of the membrane-embedded region and contribute significantly to the net resistance, in contrast to the empty nanopore where the resistance is mainly attributed to the membrane-embedded region. This likely explains why the measured nanopore current depends sensitively on the conformational states of the trapped protein. The simulation further revealed that the holo MBP conformer, which is more compact, is trapped more deeply within the ClyA lumen than the apo conformer (Table 1), which leads to greater blockage of the current. The more elongated apo MBP is trapped in the higher, larger-diameter part of the lumen, thus leaving more solvated-ion accessible regions and gives rise to smaller unit resistance than bound MBP.

**Figure 3.**
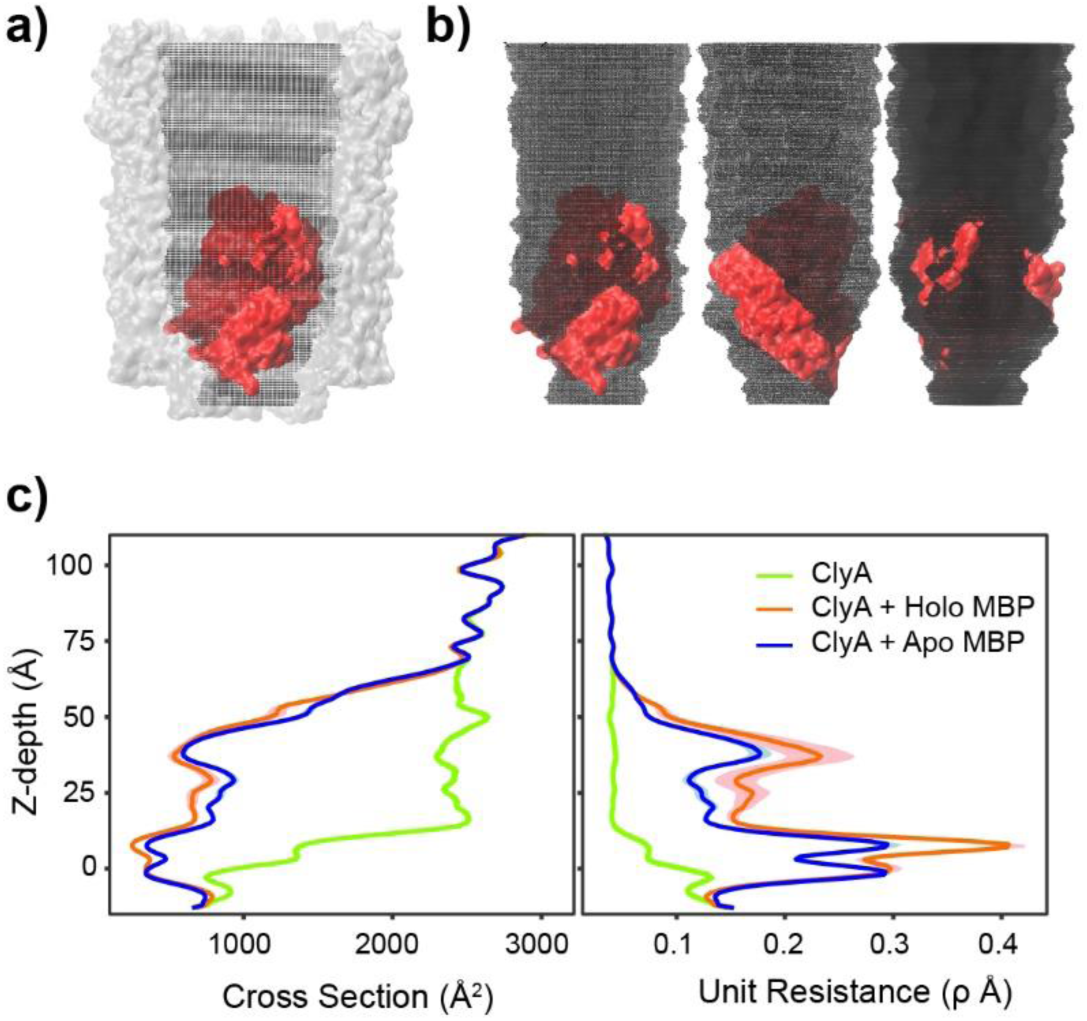
Molecular basis of ClyA nanopore current blockades (a) A representative conformation of ClyA nanopore with a trapped MBP in the apo state. The color code for ClyA and MBP are grey and red respectively. Accessible volume of solvated ions inside the pore is shown in black. (b) Alternative views of the solvated ions accessible volume. (c) Nanopore profile of solvated-ions accessible cross section and unit resistance. Results from holo and apo MBP simulations are shown in red and blue. The green line represents estimation for empty ClyA nanopore without trapped MBP. The colored shades are uncertainty estimated from the snapshots chosen periodically in the last 500 ns of the simulations. The unit of resistance is using unit resistance which is 100ρ. Here ρ is the physical parameter of charge density and assumed to be 1 in the estimation.

**Table 1.** Simulation.

| Force<br>(kcal/mol/Å) | Conformation | Length<br>(μs) | Depth<br>(Å) | Net Unit Resistance* |  |
| --- | --- | --- | --- | --- | --- |
|  |  |  |  | ClyA | ClyA + MBP |
| 0.1 | apo | 2.0 | 29.3 ± 0.3 | 6.6 ± 0.0 | 15.7 ± 0.7 |
| 0.1 | holo | 2.0 | 30.5 ± 0.7 | 6.6 ± 0.0 | 13.3 ± 0.3 |
| 0.5 | apo | 2.0 | 26.5 ± 0.4 | 6.6 ± 0.0 | 22.1 ± 0.4 |
| 0.5 | holo | 1.0 | 27.4 ± 0.5 | 6.6 ± 0.0 | 17.0 ± 0.3 |
| 1.0 | apo | 1.0 | 26.3 ± 0.2 | 6.6 ± 0.0 | 22.6 ± 0.6 |
| 1.0 | holo | 1.0 | 27.2 ± 0.3 | 6.6 ± 0.0 | 17.2 ± 0.2 |

\* The last 500 ns of the simulation were used to calculate the z-depth and the net unit resistance. The net unit resistance is integrated the unit resistance through the whole pore. The uncertainty is due to the calculation using periodical snapshots in the last 500 ns.

Once net unit resistance is converted into current blockade, models predict that ClyA with trapped holo MBP will cause a 6 % to 10 % greater current blockade than ClyA with apo MBP, depending on the net drifting force constants used (which mimic the electroosmotic force under constant probing voltage; see Methods) (Table 1). The magnitudes of current reduction upon MBP trapping as well as ligand binding predicted by the simulation are in quantitative agreement with the actual measurements (Fig. 2). The model also indicated that when MBPs are trapped more deeply with a stronger force constant, the tilting angle of holo MBP is larger than that of apo MBP (Supplementary Fig. 5). This suggests that MBP adjusts its orientation as a result of nonspecific ClyA-MBP interactions and topology. These results suggest that it may be possible to resolve additional details of the conformational changes of a trapped protein by adjusting the applied voltage.

## Anomer maltose-specific binding states

The current model of maltose-MBP binding has two states—bound and unbound. Therefore, we were surprised to find that the ClyA nanopore resistance recordings showed four states (Fig. 2b&c). One possible explanation for this result is that states 1, 2, and 3 represent a single holo MBP conformer that can become trapped within the lumen of the ClyA nanopore in three different configurations. If this was the case, the relative abundance of current states 1, 2, and 3 would be determined by the affinity constants of the MBP/ClyA complex in these alternative configurations and independent of the ligand concentration (Supplementary Fig. 6). However, our data clearly show that the relative abundance of the three states varies with ligand concentration (Fig. 2b&c).

Another explanation for the additional states is that they result from an anomeric effect. Maltose is a reducing sugar with an α-anomer and a β-anomer that coexist in solution and mutarotate via an aldehyde intermediate (Fig. 4a). A previous NMR study showed the tritium-labeled α-maltose and β-maltose bound to MBP with different affinities^24^. Inspired by this observation, we hypothesized that the multiple ligand-bound states represent MBP bound with different maltose anomers. We were not able to find a source of anomerically pure maltose to directly test the hypothesis. Instead, we introduced maltose phosphorylase into the buffer to deplete α-maltose *in situ* (Fig. 4a). Maltose phosphorylase is a 180 kDa enzyme that specifically hydrolyzes α-maltose and generates glucose and β-D-glucose-1-phosphate (Fig. 4a)^25^. The spontaneous interconversion between α-maltose and β-maltose is a slow process with a rate constant of ∼0.009 min^−1^ and a half-life of ∼ 60 min^26^. The turnover rate of 0.25 unit of maltose phosphorylase is 1.5 × 10^17^ min^−1^. Thus, the addition of maltose phosphorylase will both deplete the existing < 1 μM α-maltose in the recording chamber within one minute and prevent α-maltose from accumulating in a significant quantity. This results in a solution of primarily β-maltose^27^. Control experiments showed that neither maltose phosphorylase nor its reaction products interfered with the current recording of MBP (Supplementary Fig. 4).

After a single MBP protein was trapped inside the ClyA nanopore, we added 1 μM maltose and 10 μM sodium phosphate into the *cis* chamber and observed the typical trace with four current levels (Fig. 4b). Then 0.25 unit of maltose phosphorylase was added to the system, the current signal showed all three bound states for the first minute. After 1 min incubation the state 2 signal disappeared but state 1 and some state 3 events were still present (Fig. 4b bottom). The frequencies of state 1 and state 2 before and after the enzyme addition are summarized in Fig. 4c, with outliers removed on the basis of Grubbs’ test. Depletion of α-maltose led to the frequency of state 2 dropping from 0.18 ± 0.17 s^−1^ to 0 (n=4), while the frequency of state 1 increased from 4.28 ± 0.57 s^−1^ to 5.59 ± 0.89 s^−1^. Because depletion of α-maltose led to a reduction in the state 2 signal, the data strongly suggest that state 2 represents MBP bound to the α-maltose anomer. In addition, we would expect β-maltose to bind to MBP more frequently after the α-maltose was depleted by the enzyme, because of the reduced competition from α-maltose, and indeed the frequency of state 1 did increase after the α-maltose was depleted. For these reasons, we assigned state 1 as the β-maltose-bound state of MBP.

**Figure 4.**
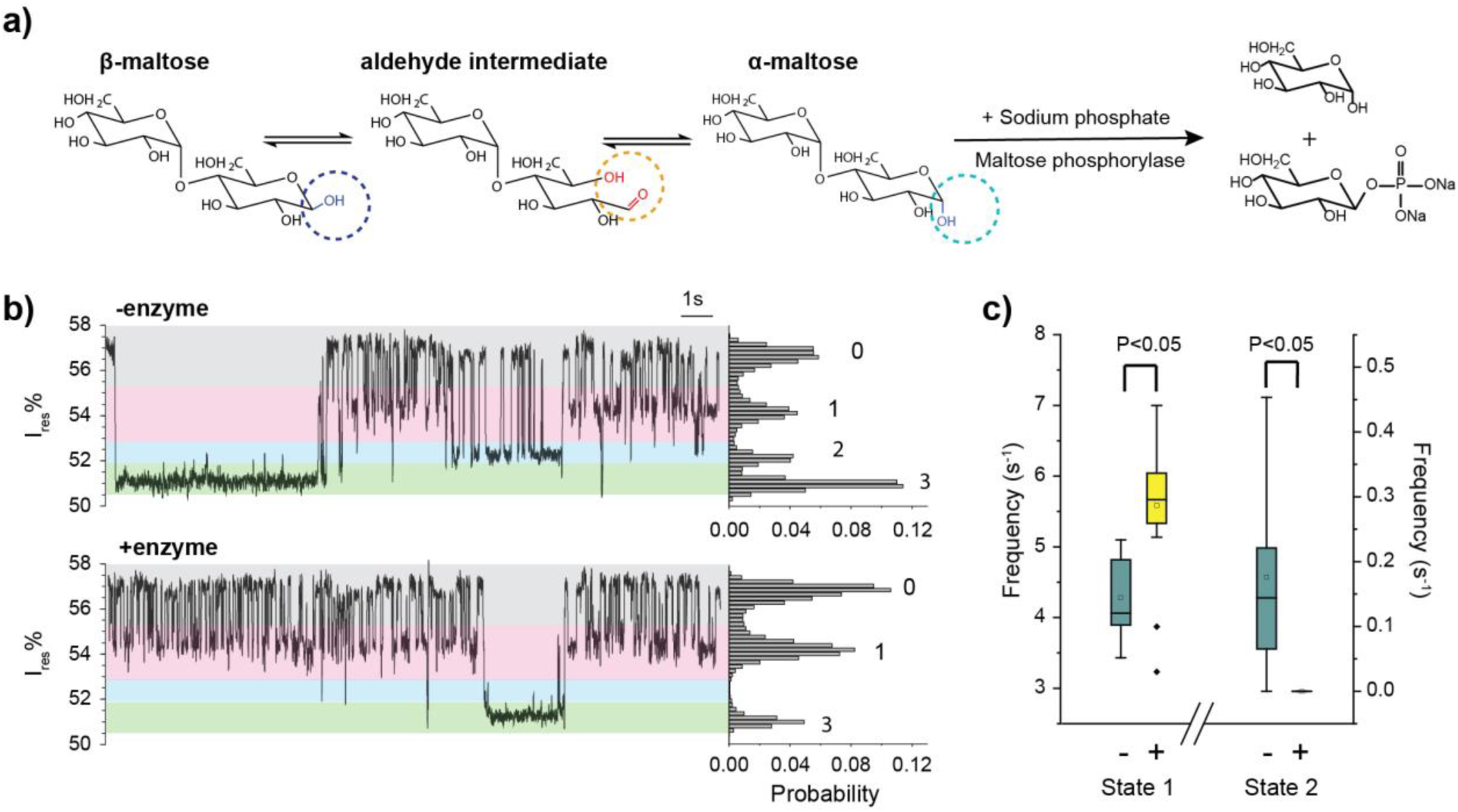
Identification of anomeric binding to MBP by depletion of α-maltose with ClyA. (a) Reaction schemes of maltose mutarotation and maltose phosphorylase-catalyzed phosphorolysis of α-maltose. (b) Representative current traces and the corresponding all events histograms of a ClyA pore with a trapped MBP before and after removal of α-maltose bymaltose phosphorylase. (c) Frequency of state 1and 2 before and after depletion of α-maltose. The two-tailed t-test was performed and the confidence interval is 95 %. Data were collected from 9 individual events before the depletion of α-maltose and 20 trapped MBP events after the depletion. All the measurements were performed in 150 mM NaCl, 15 mM Tris-HCl, pH 7.5 at −80 mV by applying a Bessel low-pass filter with a 2 kHz cutoff and sampled at 5 kHz.

State 3 events were rare, with a frequency of 0.05 ± 0.04 s^−1^ before the depletion of α-maltose and 0.01 ± 0.02 s^−1^ after the depletion of α-maltose. Because of the large uncertainty in these measurements, we cannot accurately assess the change in the abundance of state 3 before and after enzyme addition. Two possibilities were considered for the identities of state 3. One explanation is that state 3 results from MBP bound to an aldehyde intermediate. An equilibrated maltose solution contains 42 % α-maltose and 58 % β-maltose^28^, as well as a very small fraction (0.0026 %) of aldehyde intermediate. ^29^ The frequency of state 3 is significantly lower than that of the other two states, and thus it is possible that state 3 represents MBP bound to the aldehyde intermediate. Another possibility is that state 3 represents a second conformation of β-maltose bound to MBP. An NMR study of tritium-labeled maltose showed two separate chemical shifts for β-maltose upon interaction with MBP, which suggests that β-maltose can bind to MBP in two different modes^24^. We investigated these two possibilities through the analysis of thermodynamic and kinetic parameters as described in the next section.

To further confirm that the three bound states were related to the anomeric forms of the ligand, we tested MBP bound with β-cyclodextrin, a ligand that does not undergo mutarotation because of its cyclic-ring structure^30^. In the presence of β-cyclodextrin, we observed one major current level similar to state 1 of maltose. This indicates that β-cyclodextrin has only one binding mode with MBP (Supplementary Fig. 4). In addition, we observed that MBP bound to maltotetraose, another reducing ligand, exhibited three ligand-bound current levels similar to those of maltose and maltotriose (Supplementary Fig. 4). Taken together, these data strongly indicate that the multiple current levels of ligand-bound MBP are derived from its interaction with the α- and β-anomers of the reducing maltooligosaccharides.

## Kinetic and thermodynamic analysis of MBP-ligand interactions

The ability of the nanopore tweezer to monitor the electrical current fluctuation for extended periods has allowed us to derive precise kinetic and thermodynamic parameters of MBP-ligand interactions. We first titrated trapped MBP with increasing concentrations of maltose or maltotriose (Table S1) and calculated the dissociation constant (K_d_) of the ligand-MBP bound by using accumulative abundance of the three bound states (Table 2). The dissociation constants calculated by this method were 0.27 ± 0.04 μM (Fig. 5a) for maltose, which is similar to the 0.75 ± 0.13 μM measured by isothermal titration calorimetry in our lab (Supplementary Fig. 7) as well as reported values of 0.35 to 1.2 µM measured by fluorescence titration and isothermal titration calorimetry^23,31^. The calculated K_d_, 0.15 ± 0.01 μM (Fig. 5b), for maltotriose is comparable to the previously determined value of 0.16 to 0.7 μM^31^. These nanopore-derived values are consistent with previous findings^31^ showing that maltotriose has a higher affinity for MBP than maltose.

**Figure 5.**
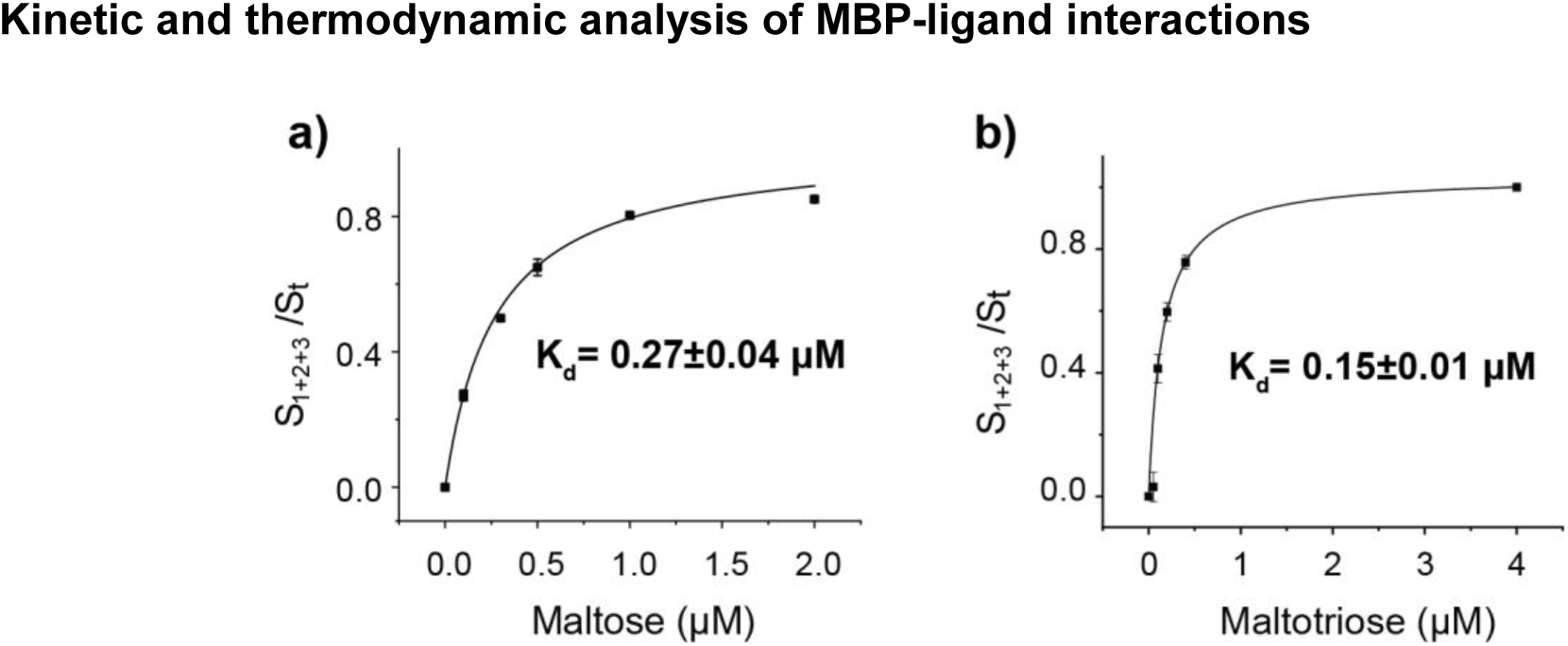
Titration curves and binding affinities of MBP for maltose (a) and maltotriose (b).

**Table 2.** Kinetics and thermodynamics of MBP bound to different ligands.

| Ligand | State | Anomer | $K_d$ ($\mu\text{M}$) | $k_{\text{off}}$ ($\text{s}^{-1}$) | $k_{\text{on}}$ ($\mu\text{M}^{-1}\text{s}^{-1}$) |
| --- | --- | --- | --- | --- | --- |
| Maltose | 1 | $\beta$ | $0.31 \pm 0.01$ | $13.3 \pm 1.6$ | $43.6 \pm 0.1$ |
| | 2 | $\alpha$ | $0.54 \pm 0.05$ | $3.5 \pm 0.2$ | $6.5 \pm 0.1$ |
| | 3 | $\beta$ | $0.42 \pm 0.02$ | $1.3 \pm 0.5$ | $3.1 \pm 0.4$ |
| Maltotriose | 1 | $\beta$ | $0.12 \pm 0.00$ | $4.8 \pm 0.3$ | $40.8 \pm 0.1$ |
| | 2 | $\alpha$ | $0.24 \pm 0.01$ | $1.6 \pm 0.1$ | $6.6 \pm 0.1$ |
| | 3 | $\beta$ | $0.38 \pm 0.02$ | $1.9 \pm 0.4$ | $5.0 \pm 0.2$ |

The errors represent the standard deviations of 5 independent pores and 51 trapped MBPs.

Because the three bound states were well resolved, we were also able to calculate the K_d_ for each anomeric species (Supplementary Fig. 8). The high temporal resolution of the nanopore-based current recording technique made it possible to capture transitions between states to derive the kinetic parameters. The dissociation rate constant, *k*_*off*_, was derived from the dwell time (τ_off_) of each binding model, which is independent of ligand concentration (Supplementary Fig. 9).

Quantification of kinetic parameters have also shed light on the identity of state 3. For this, we compared kinetic parameters calculated with two different assumptions, first, that state 3 represents MBP bound to an aldehyde intermediate of the sugar molecule, and second, that state 3 represents an alternate conformation of MBP bound to the β-anomer of the sugar molecule. Under the first assumption, we derived a *k*_*on*_ of 6.9 ± 2.8 × 10^10^ M^−1^s^−1^, *k*_*off*_ of 1.3 ± 0.5 s^−1^, and *K*_*d*_ of 1.9 ± 0.1 × 10^−5^ μM. The association rate calculated with this assumption was four orders of magnitude higher than that of the other two isomers and exceeds the known diffusion limit for any enzyme^32–35^. This suggests that state 3 is highly unlikely to represent MBP bound to the aldehyde species. As such, we believe that state 3 may represent a second conformation of MBP bound to the β-anomer of maltose. Based on this assignment, the thermodynamic and kinetic parameters of all three states, are summarized in Table 2. These data show that the β-anomer exhibits the highest affinity for MBP when it adopts binding mode 1. Maltose has a higher K_d_ than maltotriose at each binding mode. Maltose and maltotriose have similar association rates but different dissociation rates, which results in their different binding affinities.

## Conclusions

We applied the ClyA nanopore to capture a single MBP molecule and study its interaction with ligands over multiple cycles of association and dissociation. We observed that MBP could adopt three distinct binding modes upon interacting with ligands. Combining biochemical experiments and kinetic and thermodynamic analyses, we show that these three ligand-bound modes are induced by MBP binding to different anomeric forms of maltooligosaccharides. So far, X-ray crystallography obtained MBP structures with the preferential binding of only the α-form of linear maltooligosaccharides^23,36^, despite ^3^H NMR studies with tritium-labeled ligands showed that MBP could bind to both α- and β-anomers^24^. To our knowledge, it is the first time that MBP has been observed to adopt different conformations upon binding to anomeric-sugars. Nevertheless, the results can be explained by the structural arrangement of the substrate binding site within MBP. Regardless of the length of the oligosaccharides, the reducing glucose units of all oligosaccharides are buried in identical binding pockets that feature two amino acids, Asp 14 and Lys 15, which form hydrogen bonds with the α-anomeric hydroxyl^23^. The interactions between an β-maltooligosaccharide and Asp 14/Lys 15 would be disrupted because the β-anomic hydroxyl flips from the axial to the equatorial position. The loss of the two hydrogen bonds may cause the β-anomer–bound MBP to adopt a slightly different conformation than the α-anomer–bound MBP. The resulting difference in conformation of the sugar-MBP complex could be too small to be resolved by NMR^37^; yet it generated distinct current levels measurable by the ClyA nanopore tweezer. Interestingly, the technique even detected two types of β-maltooligosaccharide bound states, supporting the previous observation by tritium NMR spectroscopic studies^49–51^.

Computer modeling and simulations have provided the molecular basis for the sensitivity of the ClyA trapping measurement to resolve conformations of the trapped MBP. The trapped protein in the lumen gives rise to additional constrictions above the membrane-embedded region, which can depend sensitively to the protein conformation (and configuration of entrapment). Specifically, the holo MBP, which is more compact than apo MBP, becomes trapped at a deeper location within the lumen of the ClyA nanopore and therefore causes a greater current blockade than apo MBP does. However, the present coarse-gained model did not reveal details about conformational differences between MBP bound with α-anomers and β-anomers. In the future, atomistic simulations would be useful to interpret the distinct current levels triggered by MBP bound to different anomers of the ligand. Particularly, the MD simulation of MBP may provide evidence for the two possible binding modes of β-anomers observed here and in the previous NMR study.

A major advantage of nanopore analysis of protein functional motion is that the time-resolved current fluctuations can be used to quantify thermodynamic and kinetic properties of the conformational transitions. Binding affinities of maltose and maltotriose for MBP calculated from nanopore measurements are highly consistent with reported values from multiple bulk assays including our isothermal titration calorimetry experiments. The kinetic rate constant *k*_*on*_ *and k*_*off*_ calculated from the ClyA experiments approximate the results obtained from the rapid-mixing, stopped flow technique^38^. Our calculated values indicate that maltotriose has a higher affinity for MBP than maltose does, which is consistent with previous structural and biochemical analyses^23,31^. Our data also show that within the similar binding mode, the *k*_*on*_ values of maltose and maltotriose are very similar, while the *k*_*off*_ of maltose is three times higher than that of maltotriose. These data could be readily explained by the structures of MBP because the additional glucose unit of maltotriose could make extra contact with the aromatic residues within the binding pocket of MBP.

The single-molecule ClyA nanopore tweezer study has further elucidated the anomic specificity of reduced maltooligosaccharides for MBP, a phenomenon which would be difficult to evaluate with other techniques because mutarotation occurs spontaneously in solution. Our results showed that α-maltose and α-maltotriose had lower *k*_*off*_ values for MBP than β-maltose and β-maltotriose. This result is consistent with the structural arrangement of the ligand binding pocket that allows the α-anomer form two more hydrogen bonds with MBP than the β-anomer. Interestingly, the β-maltodextrins, exhibited significant faster association rates than the α-anomers for both maltose and maltotriose in our measurements. As a result, MBP-bound β-anomer has a slightly higher affinity than α-anomer in the predominant binding modes, state 1 and state 2 (Table 2). This observation is contradictory to results obtained from tritium NMR spectroscopy that show that α-maltose has a ∼2.5-fold higher affinity for MBP than β-maltose^24^. Here we observe that both the apo state 1 and the β-maltose bound state 1 has the same current level, which suggests that the two states have similar structures. Thus, we surmise that the higher affinity of the β-anomer observed here may be because the trapped apo-MBP adopts conformational states that favor β-anomer association.

The ClyA trapping method deviates from classic nanopore sensing techniques in which analytes interact transiently with or translocate through nanopores^39^. In this work, the funnel-shaped ClyA was used as a molecular tweezer to trap the analyte MBP for a duration (average ∼200 s) long enough to observe the action of MBP over multiple cycles of ligand interactions. The label-free, single-molecule nanopore tweezer allows continuous monitoring of the protein’s motion at 10 μs to ms resolution. The approach could be modified to manipulate the trapping time by adjusting the applied potential. In addition, multiple analyte molecules could be probed separately by simply reversing the voltage to eject the trapped molecule and capture a new one. When combined with state-of-art multiple channel recording systems^40^, the method is highly scalable and has the potential to become a high-throughput approach for monitoring protein-ligand interactions. Despite the advantages, the nanopore sensing technique described here has some limitations. Unlike the FRET and NMR, current signals do not contain distance information of the states that can be used for structure calculations. Therefore, assigning the current states to biologically relevant conformational states can be difficult without previous knowledge of the conformers. To overcome the limitation, computational simulations will be essential to interpret the time-series of state transitions revealed by nanopore trapping measurements.

In summary, this study establishes the viability of applying the ClyA nanopore tweezer to investigate the conformational dynamics of MBP by continuously monitoring its motion with high temporal and spatial resolution. We believe that this approach will complement conventional smFRET and NMR methods and provide a valuable tool for comprehensive analysis of protein folding, enzymatic catalysis, and biomolecule recognition, as well as drug screening.

## Methods

### Protein preparation

The ClyA gene with C-terminus His6 was cloned into a pT7 plasmid vector using *HindIII* and *Xhol* restriction digestion sites. The MBP gene with N-terminus His8 was cloned into a pT7 plasmid vector using *HindIII* and *NdeI* restriction digestion sites. ClyA monomer and MBP were expressed and purified with the similar protocol according the previous study^19^. Both plasmids were transformed into BL21(DE3) Competent *E. coli* cells to express proteins. Proteins were purified by using Ni-NTA affinity chromatography. The purified proteins were further purified by FPLC to remove the potential aggregates. Protein concentration was then determined by BCA assay and stored in −80 °C freezer for future use. The ClyA monomer was assembled into oligomer with 0.2 % DDM at room temperature for 20 min and then stored in 4 °C fridge for electrophysiology experiments.

### Single channel recording experiments with planar bilayer

Single channel recording is similar as described before^41^. Briefly, a ClyA nanopore was inserted into a DphPC lipid bilayer separating two chambers. Both chambers were filled with 900 µL buffer (150mM NaCl, 15 mM Tris-HCl, pH 7.5). ClyA was added from the *cis* chamber and inserted into the bilayer spontaneously. MBP was also added from the same chamber to be trapped in nanopore. −80 mV potential was applied across the bilayer and ionic current through nanopore was monitored in voltage-clamp mode by integrated patch clamp amplifier (Axopatch 200B, Molecular Devices). The signal was acquired by an analog-to-digital converter (Digidata 1440A, Molecular Devices) at a sampling rate of 50 kHz after processing with a 4-pole lowpass Bessel filter at 2 kHz. Data was recorded by Clampex software (Molecular Devices). Experiments were conducted at 23 °C.

### Simulation System setups

Due to the large size, the system was represented using the MARTINI coarse-grained (CG) force field to allow sufficient sampling of potentially heterogeneous and nonspecific interactions between MBP and ClyA. MARTINI represents an approximate 4 to 1 mapping of heavy atoms to CG beads^42–44^. Each CG bead is assigned a specific particle type based on the chemical nature of the underlying structure. The non-bonded interactions have been parameterized to capture to experimental thermodynamic data^42,43^. Importantly, despite the CG nature, the MARTINI force field has been extensively parameterized provide a quantitative description of the thermodynamic balance of protein-water, protein-protein and water-water interactions. It has been successfully used to describe protein-protein interactions in both aqueous and membrane environments^45,46^. Therefore, the MARTINI force field is expected to be appropriate for probing the interactions between MBP and ClyA.

Unbound MBP has a wider angle between two domains. The Cα RMSD between ligand bound and unbound MBP structures is 3.9 Å. The ClyA nanopore has 12 units of protomer, and each of them has 285 amino acid residues. The atomistic structures were mapped onto the MARTINI CG representation separately using CHARMM-GUI ^47^. MARTINI requires additional structural restrains to remain the protein structure. This was achieved by elastic network restraints with a force constant of 1.5 kcal/mol/Å2. The force constant was assigned by matching the flexibility of CG simulation to results from atomistic runs. For this, pioneer atomistic and CG simulations were performed to validate the force constant of restraining secondary structure in MARTINI CG model. In atomistic simulation, MBP was solvated in a cubic water box of 84.76 Å with TIP3P water^48^. CHARMM was used to perform Langevin dynamics using CAHRMM 36m force field in constant temperature 298 K and constant pressure 1.0 atm^49,50^. The SHAKE algorithm was applied to all hydrogen-containing bonds to allow a 2.0 fs timestep^51^. Particle mesh Ewald was utilized for electrostatics with a real-space cutoff of 13 Å^52^. Van der Waals interactions were cut off at 13 Å with a switching function between 12 and 13 Å. In CG simulation, MBP was solvated with MARTINI water box of 126.40 Å by 140.64 Å by 152.66 Å. NAMD 2.12 was used to perform the CG simulation with elastic network restraint (1.5 kcal/mol/Å2) applied to MBP. Parameters used are 5 fs timestep, switch distance cutoff 9.0 Å, real-space cutoff 12.0 Å, dielectric constant 15, and temperature 300 K. The martiniSwitching was on to enable the MARTINI Lennard-Jones switching function. The root-mean-square fluctuation (RMSF) profiles of MBP calculated from the last half of 50 ns atomistic and CG simulations is shown in Supplementary Fig. 5b. The RMSF comparison validates that the coarse-grained model is much stable than atomistic model for nonstructural region, but for the structural region, the CG model remains the same properties as atomistic simulation. That indicate the elastic network restraints in CG model do remain the protein structure property as atomistic model. The geometry of ClyA nanopore is well-characterized, and thus cylinder boundary condition with radius 62 Å and length 160 Å was used to reduce the computational cost for further production simulations. During the whole course of simulation, the MD parameters were set as described in the pioneer CG simulation. Additionally, position restraint with force constant 5 kcal/mol/Å^2^ was applied to MARTINI backbone atoms of ClyA nanopore to restraint the pore shape.

### Generation of initial configurations and production simulations

To generate initial conformations with MBP captured inside the nanopore, MBP was first placed 30 Å above the pore entrance, and then 10 ns constant velocity pulling simulations were performed to guide MBP enter the pore. The pulling velocity was 0.00005 Å/timestep (i.e., 10 Å/ns) applied on the center of mass of MBP with direction to the bottom of the pore. Given the elongated geometry of MBP and the pore size, MBP can enter the ClyA pore in only two possible orientations, the N-terminus of MBP facing either up or down with the alignment to its the principal axis (Supplementary Fig. 5a shows the N-terminus up). The dipole moments of proteins were calculated using the PDB structures with standard CHARMM 36m atomic charges. The dipole-dipole interaction potentially provides a bias favoring the N-terminus up orientation by ∼0.26 kcal/mol in aqueous solution. As such, the simulations have focused exclusively on cases where MBP is captured in the N-terminus up orientation. After the MBP entered the nanopore, 60 ns constant force pulling simulations (50 kcal/mol/Å) were performed to allow MBP travel deeper in the pore. The final coordinate of MBP inside the pore was then used as initial structure for production simulations. The initial z-depths of bound and unbound MBP are 26.6 Å and 25.9 Å, respectively. In production simulations, weak constant force pulling simulations (0.1, 0.5 and 1.0 kcal/mol/Å) were constantly performed to mimic the constant flow in the experiments. As MBP trapping free energy is about −12.6 kcal/mol from experiment, the proper force constant is arguable to be in the range of 0.1∼0.2 kcal/mol/Å.

### Data analyses

VMD was used for the structural visualizations and data analyses^53^. The z-depth is measuring the center of mass of MBP, and the origin is at the center of member. Euler angles (φ, θ, and Ψ) of MBP is using geometrical definition to identify the orientation in space. Due the pore symmetric, only the Euler angles θ needs to be considered and that reflects the tilting angle between MBP and nanopore. Both z-depth and Euler angles θ were then used as reaction coordinate for visualizing the free energy surface of sampling (Supplementary Fig. 5c). To estimate the solvated-ion cross section and unit resistance, the in-house python scripts were used. A box of gird points with 1 Å spacing was first built to cover the whole pore. The overlapped grids to the ClyA and MBP were then removed based on the radii of CG beads. We further considered chloride ion with an additional water probe as the solvated-ion with a radius of 3.2 Å to filter out the non-accessible grids. Finally, a filter was used to ensure the continuity of girds that contribute to conductance. The pore profiles of the solvated-ion cross section were then estimated based on the remaining grids (i.e., one grid ∼ 1 Å^2^ cross section). Followed by the resistance approximation, the solvated-ion cross section can be converted to unit resistance, and the net resistance can be integrated through the whole pore^54^:

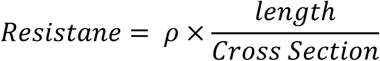

Here, length is the Z-depth of nanopore and for our estimation can be treated as 1 because the nanopore is the same. One assumption made here is charge density (ρ) doesn’t vary for apo-MBP and holo-MBP when they were trapped in nanopore. Thus, the unit resistance (ρÅ) has the reduced unit with ignoring the effect of the charge density. We equally chose 20 frames from the last 500 ns to get the statistic error of unit resistance estimation. Other analyses and plots were done with in-house scripts in R^55^.

