## Supplementary figures and table for "A ClyA nanopore tweezer for analysis of functional states of protein-ligand interactions"

### Supplementary information figures

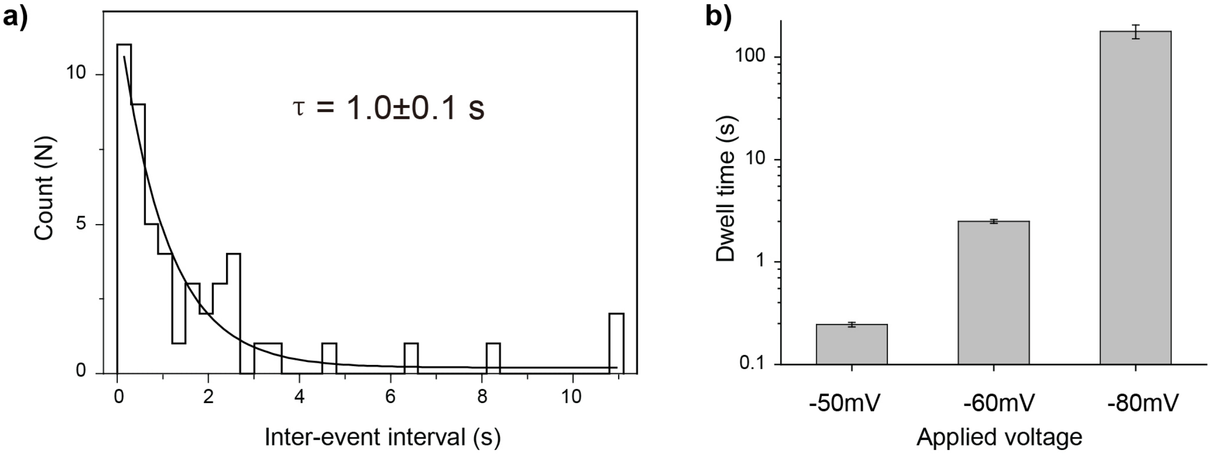

Supplementary Figure 1. Kinetic analysis of MBP trapped in ClyA. (a) The histogram shows the inter-event interval of 56 nM MBP interacting with the nanopore under -80 mV (n= 49) and fitted with standard exponential function. (b) The dwell time of 56 nM MBP interacting with the nanopore under -50 mV (n= 96), -60 mV (n= 45) and -80 mV (n= 49)

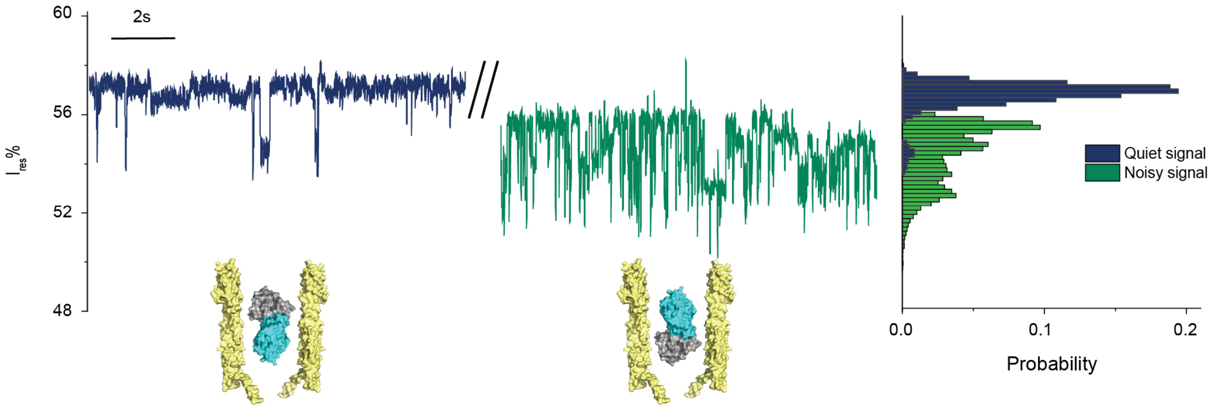

Supplementary Figure 2. MBP trapped in ClyA. Two electrical patterns observed for MBP trapped in ClyA nanopore. There was ~ 60 % chance to see the quiet signal (blue), which responded to the ligand binding and were selected in the following analysis. The noisy signal (green) has a smaller $I_{res}$ % and represents the pattern that didn't respond to the ligand binding. The histogram on the right shows the corresponding histogram of the two trapping patterns.

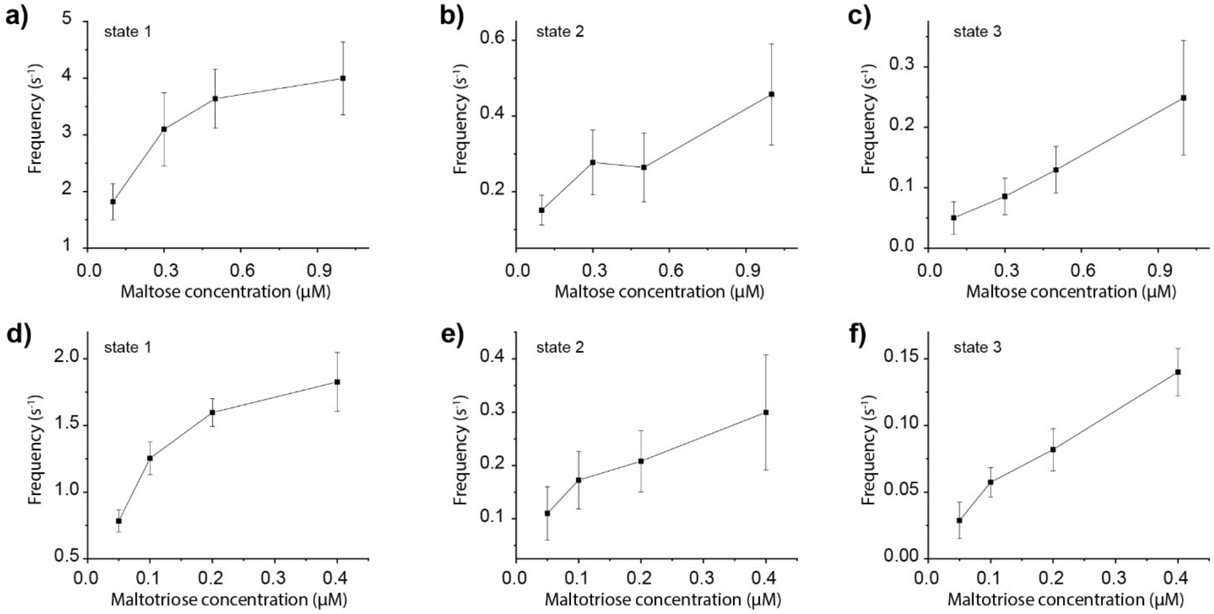

Supplementary Figure 3. Frequencies of the three MBP-bound states in varied concentrations of maltose or maltotriose. The state frequencies collected from multiple MBP/maltose and MBP/maltotriose trapping experiments were shown in (a-c) and (d-f) separately as the average SD. All the measurements were performed in 150 mM NaCl, 15 mM Tris-HCl, pH 7.5 with -80 mV applied in *trans*. Detailed information was listed in Supplementary Table 1.

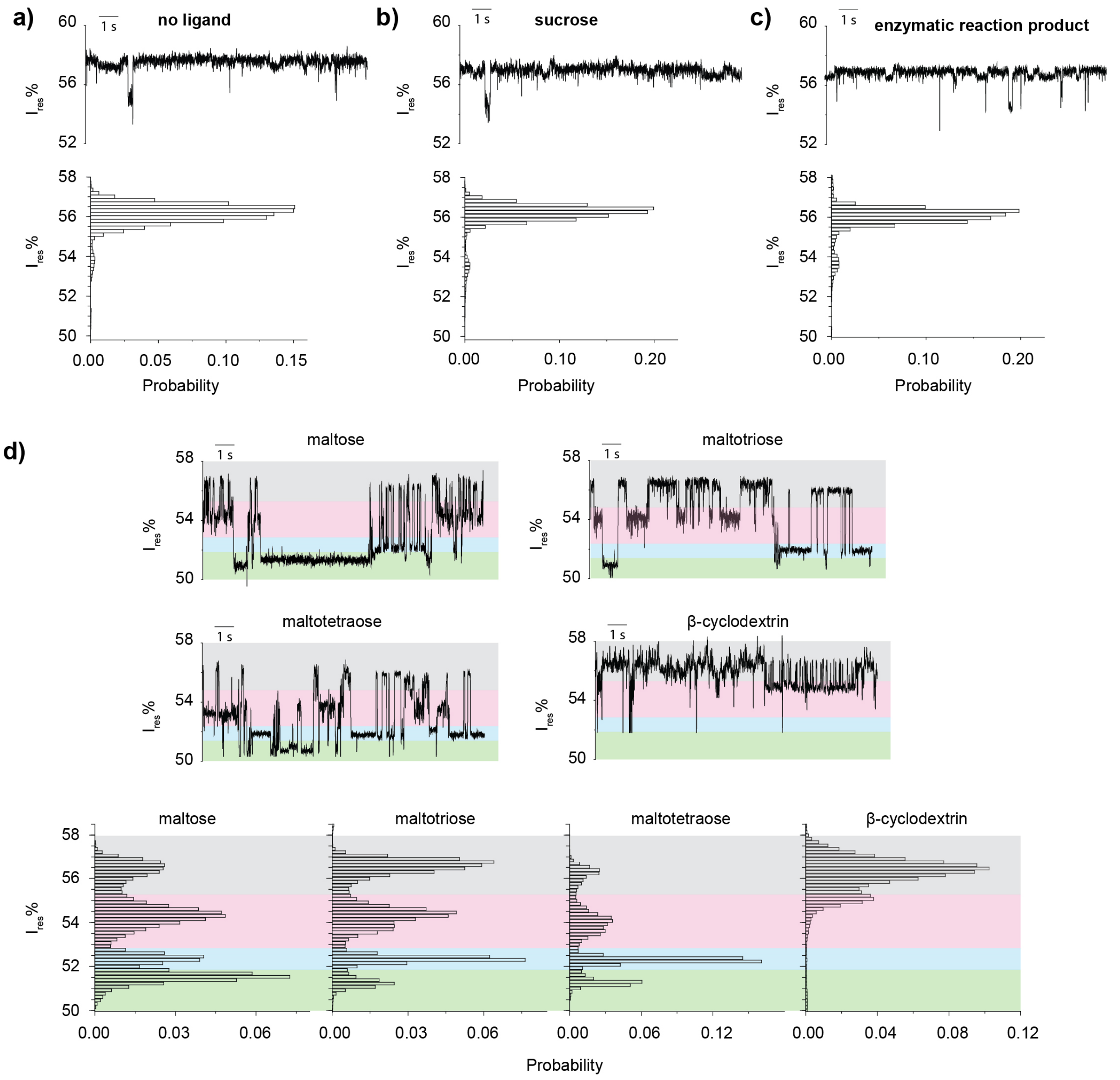

Supplementary Figure 4. Detection of MBP with non-substrates and substrates by nanopore. (a) Representative trace and blockades histogram of individual MBP confined inside ClyA. (b) Typical trace and blockades histogram of MBP with 1 µM sucrose added in the *cis* chamber measured by ClyA. (c) Representative trace and blockades histogram of MBP after addition of enzymatic reaction product with ClyA. 2 mM maltose and 2 mM sodium phosphate were incubated with 1 unit of maltose phosphorylase overnight at room temperature to obtain the concentrated reaction product. The reaction mixture (5 µl) was then diluted by 200-fold by cis buffer (150 mM NaCl, 15 mM Tris-HCl, pH 7.5). (d) Typical trace and blockades histograms of MBP with 1 µM maltose or 0.2 µM maltotriose or 0.5 µM maltotetraose or 2.5 µM β-cyclodextrin added in the *cis* chamber detected by ClyA. The current was colored into 4 states according to the state assignment in MBP/maltose interactions. All measurements were performed n 150 mM NaCl, 15 mM Tris-HCl, pH 7.5 by applying a Bessel low-pass filter with a 2 kHz cutoff and sampled at 5 kHz.

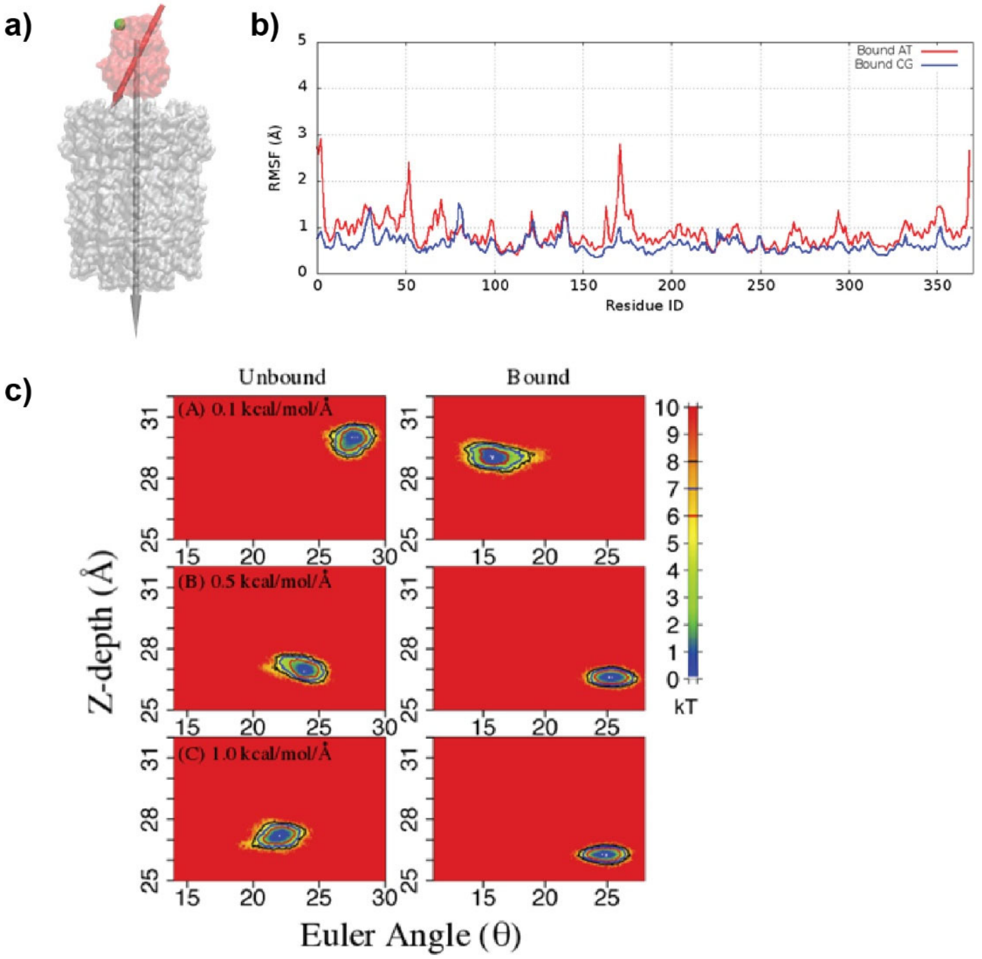

Supplementary Figure 5. Simulation. (a) Orientation of electric dipoles of ClyA and maltose-bound MBP. Both ClyA and MBP are aligned to their principle axis. The color code for ClyA and MBP are grey and red, respectively. The green sphere represents the N-terminus. In this configuration, the N-terminus is facing up. (b) The root-mean-square fluctuation (RMSF) profiles of MBP calculated from 50 ns atomistic (red) and coarse-grained (blue) simulations. (c) Free energy surface using z-depth and Euler angle θ for three force constants 0.1, 0.5, and 1.0 kcal/mol/Å. The left and right columns are from unbound and bound MBP simulations. The contour levels in red, blue and black are for 6, 7 and 8 kT respectively. The last 250 ns data of each simulation is used for data analysis.

a) Single ligand bound MBP confomer (PL) interacting with ClyA forming three complexes (PL•ClyA$_1$, PL•ClyA$_2$, PL•ClyA$_3$)

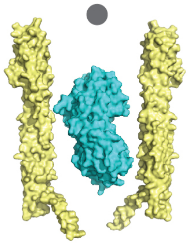

$$\mathrm{P + L \rightleftharpoons PL}$$

$$\mathrm{PL + ClyA \rightleftharpoons PL{\bullet}ClyA_1} \qquad K_{d1} = \frac{[\mathrm{PL}] \times [\mathrm{ClyA}]}{[\mathrm{PL{\bullet}ClyA_1}]}$$

$$\mathrm{PL + ClyA \rightleftharpoons PL{\bullet}ClyA_2} \qquad K_{d2} = \frac{[\mathrm{PL}] \times [\mathrm{ClyA}]}{[\mathrm{PL{\bullet}ClyA_2}]}$$

$$\mathrm{PL + ClyA \rightleftharpoons PL{\bullet}ClyA_3} \qquad K_{d3} = \frac{[\mathrm{PL}] \times [\mathrm{ClyA}]}{[\mathrm{PL{\bullet}ClyA_3}]}$$

$$[\mathrm{PL{\bullet}ClyA_1}]:[\mathrm{PL{\bullet}ClyA_2}]:[\mathrm{PL{\bullet}ClyA_3}] = \frac{1}{K_{d1}} : \frac{1}{K_{d2}} : \frac{1}{K_{d3}}$$

b) Three ligand bound MBP conformers (PL$_1$, PL$_2$, PL$_3$)

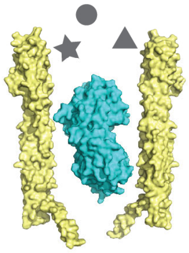

$$\mathrm{P + L_1 \rightleftharpoons PL_1} \qquad K_{d1} = \frac{[\mathrm{P}] \times [\mathrm{L_1}]}{[\mathrm{PL_1}]} = \frac{[\mathrm{P}] \times ([\mathrm{L_1}]^{\mathrm{T}} - [\mathrm{PL_1}])}{[\mathrm{PL_1}]}$$

$$\mathrm{P + L_2 \rightleftharpoons PL_2} \qquad K_{d2} = \frac{[\mathrm{P}] \times [\mathrm{L_2}]}{[\mathrm{PL_2}]} = \frac{[\mathrm{P}] \times ([\mathrm{L_2}]^{\mathrm{T}} - [\mathrm{PL_2}])}{[\mathrm{PL_2}]}$$

$$\mathrm{P + L_3 \rightleftharpoons PL_3} \qquad K_{d3} = \frac{[\mathrm{P}] \times [\mathrm{L_3}]}{[\mathrm{PL_3}]} = \frac{[\mathrm{P}] \times ([\mathrm{L_3}]^{\mathrm{T}} - [\mathrm{PL_3}])}{[\mathrm{PL_3}]}$$

$$[\mathrm{PL_1}]:[\mathrm{PL_2}]:[\mathrm{PL_3}] = \frac{[\mathrm{L_1}]^{\mathrm{T}}}{K_{d1} + [\mathrm{P}]} : \frac{[\mathrm{L_2}]^{\mathrm{T}}}{K_{d2} + [\mathrm{P}]} : \frac{[\mathrm{L_3}]^{\mathrm{T}}}{K_{d3} + [\mathrm{P}]}$$

Supplementary Figure 6. Two schemes for interpreting the multiple current levels of MBP trapped within ClyA. (a) A single ligand loaded MBP conformer (PL) interacts with ClyA in three ways to form PL•ClyA$_1$, PL•ClyA$_2$ and PL•ClyA$_3$. (b) Three different ligands interact with MBP to form three distinct conformers: PL$_1$, PL$_2$ and PL$_3$. $[L_i]^T$ represents the total concentration of each ligand in solution.

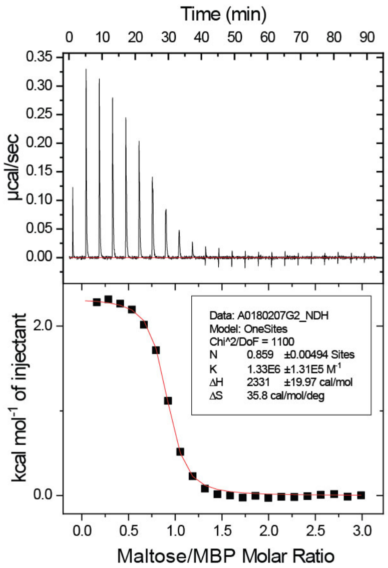

Supplementary Figure 7. Isothermal titration calorimetry assay for MBP/maltose binding affinities. Calorimetric titration curves obtained for MBP bound to maltose at 25 °C. MBP concentration was 69 µM. The titration of 1035 µM maltose included one initial injection of 0.5 µL, followed by 22 injections of 1.7 µL each. The stirring speed is 800 rpm and spacing time between injections is set to 240 s to allow the signal caused by titration to return to the baseline. Data were analyzed with the software provided by the supplier.

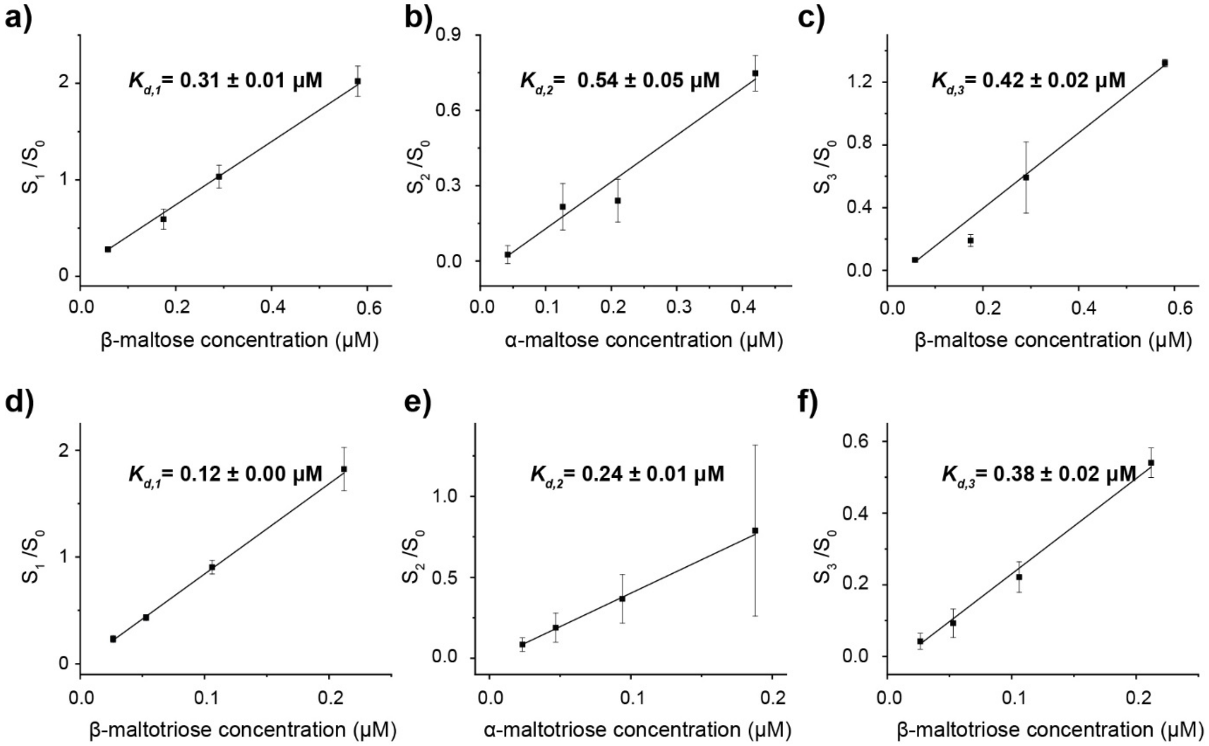

Supplementary Figure 8. The anomeric binding affinities of MBP. The binding affinities ($K_d$) for MBP bound to maltose (a-c) and maltotriose (d-f) in three binding modes were derived as described in supplementary text (determination of dissociation constant). Briefly, the binding affinity was acquired by inverting the slope of linear fitted graph of $S_i/S_0$ against the concentration of the corresponding anomer. $S_i$ indicates the area of that state and $S_0$ indicates the area of the unbound state. The concentration of α-anomer and β-anomer concentration can be obtained according to their ratio in the equilibrated maltose or maltotriose solution.

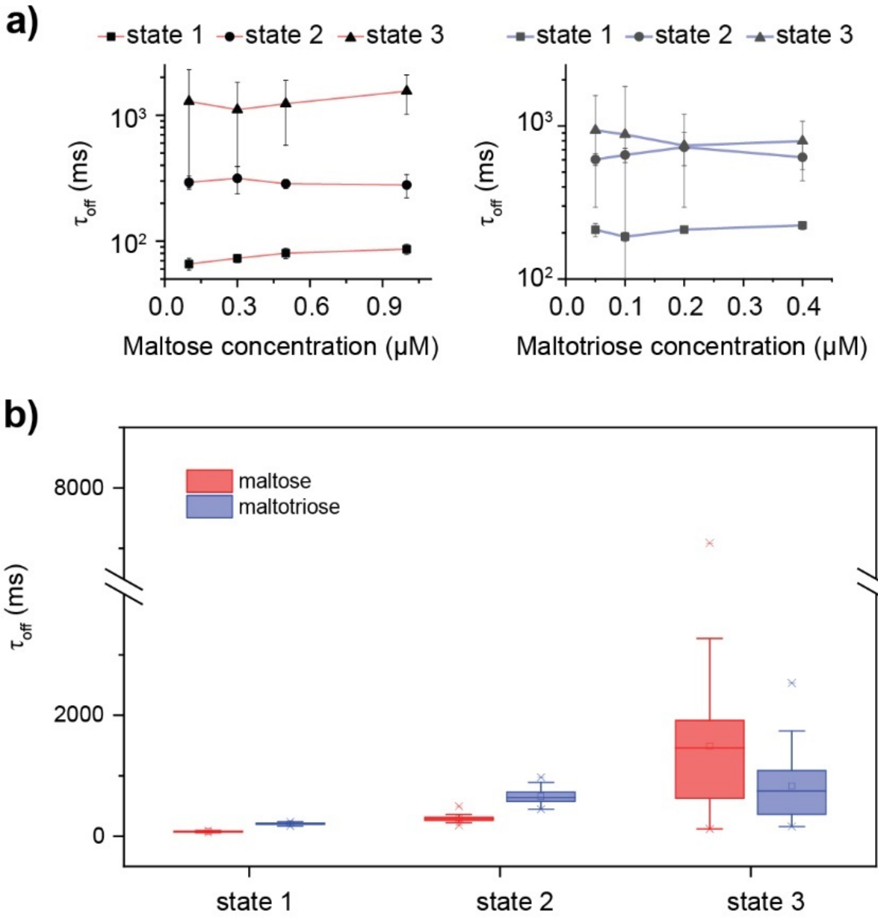

Supplementary Figure 9. Dissociation time of ligands from MBP exposed to various ligand concentrations in three binding modes. (a) The relationship between dissociation time ($\tau_{off}$) and ligand concentration in state 1 ( ¸ ), state 2 (¿ ) and state 3(s ). The MBP binding to maltose was shown in red and the binding to maltotriose was represented in blue. (b) The parallel comparison of between MBP bound with maltose and maltotriose. The dissociation time were the combined results from that binding mode regardless of the ligand concentration. The MBP binding to maltose was shown in red and the binding to maltotriose was represented in blue.

Supplementary table S1: Events collected from different MBP individuals trapped in ClyA exposed to various maltose or maltotriose concentration.

| Ligand | Concentration | Dwell Time (s) | Events of state 1 | Events of state 2 | Events of state 3 |
|---|---|---|---|---|---|
| maltose | 0.1 µM | 816 | 1691 | 63 | 15 |
| | | 928 | 1899 | 115 | 25 |
| | | 392 | 630 | 74 | 14 |
| | | 1063 | 2364 | 215 | 53 |
| | | 1164 | 1655 | 180 | 40 |
| | | 399 | 592 | 54 | 37 |
| | | 823 | 1334 | 122 | 67 |
| | | 820 | 1698 | 143 | 51 |
| | 0.3 µM | 653 | 2390 | 143 | 53 |
| | | 616 | 2328 | 160 | 48 |
| | | 183 | 691 | 26 | 4 |
| | | 910 | 2417 | 369 | 106 |
| | | 569 | 1515 | 194 | 56 |
| | | 987 | 2772 | 275 | 80 |
| | | 478 | 1628 | 168 | 56 |
| | | 488 | 998 | 107 | 45 |
| | 0.5 µM | 613 | 1762 | 71 | 45 |
| | | 179 | 600 | 55 | 25 |
| | | 660 | 2955 | 241 | 61 |
| | | 395 | 1630 | 110 | 43 |
| | | 279 | 1109 | 46 | 35 |
| | | 1000 | 3443 | 316 | 153 |
| | | 997 | 3548 | 208 | 197 |
| | | 993 | 3263 | 352 | 146 |
| | 1.0 µM | 1018 | 5525 | 647 | 430 |
| | | 817 | 3186 | 290 | 127 |
| | | 1588 | 6831 | 586 | 244 |
| | | 851 | 3204 | 335 | 155 |
| | | 454 | 1633 | 283 | 93 |
| | | 551 | 1976 | 176 | 183 |
| | | 1207 | 4093 | 712 | 351 |
| | | 1082 | 4329 | 403 | 269 |
| maltotriose | 0.05 µM | 432 | 381 | 53 | 19 |
| | | 300 | 204 | 40 | 8 |
| | | 608 | 469 | 90 | 20 |
| | | 2101 | 1686 | 78 | 24 |
| | 0.1 µM | 1314 | 1436 | 308 | 94 |

| | | | | | |
|---|---|---|---|---|---|
| | | 294 | 401 | 59 | 18 |
| | | 633 | 731 | 122 | 39 |
| | | 312 | 407 | 32 | 15 |
| | | 918 | 1243 | 121 | 41 |
| | 0.2 µM | 445 | 682 | 114 | 37 |
| | | 455 | 739 | 90 | 29 |
| | | 611 | 868 | 167 | 59 |
| | | 339 | 582 | 38 | 26 |
| | | 532 | 869 | 99 | 36 |
| | | 1052 | 1742 | 235 | 109 |
| | 0.4 µM | 339 | 513 | 156 | 56 |
| | | 780 | 1491 | 189 | 108 |
| | | 319 | 592 | 84 | 42 |
| | | 1018 | 2062 | 237 | 127 |
